# Horizontal transmission of penicillin binding protein 1A caused a nationwide spread of β-lactam resistance in pneumococci

**DOI:** 10.1101/2022.12.11.519994

**Authors:** Satoshi Nakano, Takao Fujisawa, Bin Chang, Yutaka Ito, Norikazu Kitamura, Shigeru Suga, Yasufumi Matsumura, Masaki Yamamoto, Miki Nagao, Makoto Ohnishi, Motoyuki Sugai

## Abstract

The emergence and spread of drug-resistant bacteria continue to be a global crisis. The mechanism of the resistance spread via mobile genetic elements such as plasmids is well known, however, the impact of the natural transformation on the spread remains unclear. *Streptococcus pneumoniae* is well known to be a transformable pathogen by natural competence and they become β-lactam resistance by the acquisition of chromosomal genetic elements including mutated PBPs by natural transformation. To trace the transmission of pneumococcal PBPs among nationwide pediatric population and analyze the impact of transformed PBPs to β-lactam resistance, we collected and analyzed more than 1300 isolates of *S. pneumoniae* through nationwide surveillance study for pediatric pneumococcal diseases between 2012-2017 in Japan.

We discovered a high prevalence of a specific PBP1A type (*pbp1a*-13) in β-lactam resistant pneumococci that had a 370SSMK substitution in their β-lactam binding SXXK motif, suggesting that this *pbp1a*-13 transferred horizontally between different clones resulting in emergence and spread of β-lactam resistant pneumococcal clones. Divergence dating analysis suggested that *pbp1a*-13 was inserted into major resistant lineages in the early 1990s through the 2000s, before introduction of pneumococcal conjugate vaccines in Japan. Our additional analysis for pneumococcal isolates that were recovered in the 90s in Japan suggested that *pbp1a*-13 in GPSC1 (serotype 19F-CC236) and GPSC14 (serotype 23F-CC242) isolates were the origin of the currently spread *pbp1a*-13. We provide evidence of *pbp1a* horizontal transmission at a nationwide scale and highlight the importance of PBP profile monitoring for identifying the emergence and spread of resistant pneumococci lineages.

## Introduction

*Streptococcus pneumoniae* causes various types of infectious diseases such as otitis media, pneumonia, bacteremia, and meningitis^1^. To prevent invasive pneumococcal diseases, pneumococcal conjugate vaccines (PCVs) and pneumococcal polysaccharide vaccines (PPSV) are used globally. Several countries use PCVs as one of the routine vaccines for children, while PCVs and/or PPSV are also used for elders^2^. Although the introduction of PCVs for children dramatically decreased the pathological burden^3^, pneumococcus remains one of the major pathogens associated with community-acquired pneumonia in adults^4^. Additionally, the introduction of PCVs increased the number of incidence attributable to pneumococci with non-vaccine serotype in both children and adults^5, 6, 7, 8^. Thus, *S. pneumoniae* remains an important pathogen associated with community-acquired bacterial infectious diseases globally.

While the choice of antibiotic for the treatment against *S. pneumoniae* depends on the infected organ, β-lactam antibiotics including penicillin are the first choice according to several clinical guidelines^9, 10^. Therefore, the resistance to β-lactam antibiotics is a critical issue in the treatment of pneumococcal infections, particular in the empirical therapy when clinicians have not identified the pathogen and antibiotic susceptibilities. In *S. pneumoniae*, the resistance to β-lactam antibiotics is caused by amino acid mutations within penicillin binding proteins, particular in PBP1A, PBP2B, and/or PBP2X^11^. The genes encoding these proteins are characterized by a mosaic-like structure, as *S. pneumoniae* transmit their genome to other strains via recombination events^12^. As a results, the structures of these PBPs are overall complicated while several active motifs where β-lactam antibiotics binds are conserved, such as SXXK and SXN motifs within the transpeptidase domain. In contrast, the typing method based on PBPs transpeptidase domain sequences (PBP typing), which were developed by Li Y et.al^13^, could be useful to trace the transmission origin of the “β-lactam resistance-causing PBPs” among various existing pneumococcal clones as most of the sequences are conserved and transmitted between strains. While PBP typing requires sequences of *pbp1a, pbp2b*, and *pbp2x* transpeptidase domain with more than 800 bp, recent advances in whole genome sequencing technology promote the use of the typing method in pneumococcal molecular epidemiology and aid researchers in understanding the prevalence of resistant pneumococcal lineages. However, the process responsible for progressing from an emergent lineage to nationwide spread remains unclear.

In Japan, after the introduction of PCVs in the early 2010s, meropenem resistant pneumococci increased^14, 15^. Japan is surrounded by sea and there are few population inflows from foreign counties, indicating that there are few chances of pneumococcal lineage transmission from abroad that could influence the epidemiological dynamics within Japan. Thus, we expected that this provides an adequate environment to trace the process of pneumococcal lineage expansions and/or contractions at nationwide scale. In the present study, we subjected 1305 pneumococcal isolates collected in Japan between 2012 and 2017 to whole-genome sequencing to investigate the processes associated with the emergence and spread of resistant pneumococcal lineages at nationwide scale. Subsequently, we analyzed their population dynamics to understand the characteristic of pneumococcal lineages including resistant ones.

## Materials and Methods

### Bacterial isolates

From January 2012 through December 2017, we conducted a nationwide, prospective pneumococcal infectious disease surveillance study among children in Japan^14, 16^. We collected pneumococcal isolates from children (≤15 years of age) who presented with invasive pneumococcal diseases (IPD) or non-IPD. The study design was described in detail in previous papers^14, 16^. In brief, during the study period, we collected a total of 1358 pneumococcal isolates from patients who were diagnosed with IPD or non-IPD. All isolates that grow in the stock media were subjected to whole genome sequencing.

These isolates were collected from 253 hospitals distributed in 43 out of the 47 prefectures of Japan and including 841 isolates from IPD as well as 517 isolates from non-IPD patients. Serotyping was performed using pneumococcal typing antisera (Statens Serum Institut, Copenhagen, Denmark).

### Antimicrobial susceptibility tests and the category of susceptibility

We performed antimicrobial susceptibility tests for penicillin (PCG), cefotaxime (CTX), meropenem (MEM), and erythromycin (EM) using the broth micro dilution method following the Clinical Laboratory Standards Institutes (CLSI) guidelines^17^. The 2008 CLSI standard for categorization was used to determine susceptibilities; PCG-susceptible (PG-S) and resistant (PG-R) lineages were defined based on their reaction to <0.06 and >0.06 mg/l, CTX-susceptible (CTX-S), intermediate-resistant (CTX-IR), and -resistant (CTX-R) were defined as ≤0.5, 1.0, and >1.0 mg/l, respectively, MEM-susceptible (MEM-S), intermediate-resistant (MEM-IR), and resistant (MEM-R) were defined as ≤0.25, 0.5, and >0.5 mg/l, respectively, while EM-susceptible (EM-S), intermediate-resistant (EM-IR), and resistant (EM-R) were defined as ≤0.25, 0.5, and >1.0, respectively. Multi β-lactam resistance (Mβ-lacR) was defined as an isolate resistant or intermediate-resistant to two or more drugs out of PCG, CTX, and MEM. Multidrug resistance (MDR) was defined as an isolate exhibiting Mβ-lacR with EM-IR or -R.

### Whole-genome sequencing protocol

Total genomic DNA was extracted and sequence libraries were prepared using a QIAamp DNA Mini Kit (QIAGEN, Hilden, Germany) and a Nextera XT DNA Library Preparation Kit (Illumina, San Diego, CA, USA). We multiplexed and sequenced the samples using an Illumina Nextseq system for 300 cycles (2 × 150-bp paired-end). A part of isolates was sequenced in previous studies using an Illumina MiSeq for 600 cycles (2 × 300-bp paired-end). We used Illumina HiSeq X Five (2 × 150-bp paired-end) to sequence additional 57 isolates that were collected in 1996.

### Genome assembly and annotation

After the trimming of adapters and low-quality reads using fastp v0.20.1, reads of each isolate were assembled using SPAdes v3.14.1^18^ with a k-mer size of 21, 33, 45, 57, 69, 81, 93, 105, 117, and 127. The assembly was evaluated using Quast v5.0.2^19^ The obtained draft assemblies were annotated using Prokka v 1.14.6^20^ with default parameters. Isolates with a number of coding sequence (CDS) lower than 1900 or higher than 2350 or with an N50 value lower than 30,000 were excluded from subsequent analyses.

### Sequence typing, PBP typing, and resistant and Pili gene detection

We conducted in silico MLST using mlst v2.19.0 (https://github.com/tseemann/mlst)^21^. To perform PBP typing, we extracted *pbp1a, pbp2b*, and *pbp2x* region sequences using blast+ v2.6.0^22^ and samtools v1.9^23^. We downloaded the reference fasta files for PBP typing from the CDC Streptococcus Laboratory webpage (https://www.cdc.gov/streplab/pneumococcus/mic.html) and assigned PBP type numbers to each isolate. We detected the presence of *ermB, ermTR, mefA, mefE, tetM, tetO, folA* mutation (100L and 92R), *folP* insertion, *rrgA-1* (pili1), and *pitB-1* (pili2) from the assembled contigs using blast+ v2.6. We set cut-off values to detect resistance genes as previously described by Metcalf et al.^24^.

### Phylogenetic recombination and divergence dating analysis

First, we extracted core genome of all tested isolates using the output of Prokka with roary v3.13.0^25^. We constructed a maximum likelihood tree for a core single nucleotide polymorphism (SNP) sequence using raxml-ng v0.9.0^26^ with a GTR+G4+ASC_LEWIS model. The model was decided based on the ModelTest-ng v0.1.7 results.

Second, we performed Global Pneumococcal Sequencing (GPS) clustering using Pathogenwatch (https://pathogen.watch/). Further, we identified recombination regions 27 in each cluster that included more than nineteen isolates using Gubbins version 3.1.3 with a model fitting option for Raxml-ng and 100 times of boot-strapping option. To prepare the input files for Gubbins, we obtained complete genomes for representative isolates in each cluster through hybrid assembling. The additional long read sequences were obtained using Oxford Nanopore technology. We used R9.4.1 Flow Cell and performed base call using Guppy v5.0.16 with a super-accuracy model. Hybrid-assembly was performed using Unicycler v0.4.8 with default settings or Flye v2.9 with ten times polishings through Pilon v1.23.

To address the date when PBP1A were transmitted into resistant clusters, we obtained the divergence dates using the Bayesian Markov chain Monte Carlo framework. SNP alignments without recombination regions were used as the input dataset for BEAST v1.10.4^28^. The model was selected through comparisons of the marginal likelihood using path sampling and stepping stone-based marginal likelihood estimation for a strict clock and an uncorrelated relaxed clock in a molecular clock model as well as a constant size and Bayesian Skygrid in a tree prior model. We set the MCMC lengths to obtain effective sample sizes (ESS) greater than 200 for all factors.

### Population genetic analysis

Genetic diversity indices and neutrality tests for each GPS cluster (GPSC) including 10 or more isolates were calculated using DnaSP v6.12.03^29^ To address sampling bias, we performed the analysis using only IPD isolates because the number of hospitals where non-IPD isolates were collected was much smaller than the number of those where IPD isolates were collected. We used recombination region censored nucleotide alignments for input data that were created using Gubbins. For the GPSC106 analysis, we divided the isolates into two subclusters based on a phylogenetic tree created by Gubbins because the input data were too large to analyze using DnaSP.

### Recombination region comparison

To support the hypothesis that *pbp1a*-13 horizontally transmit through recombination events, we analyzed isolates with *pbp1a*-13 that did not generate clusters in the core genome tree. First, we mapped trimmed short reads of isolates with *pbp1a*-13 to a *pbp1a*-including-recombination of the target isolate. Following identification of a set of two isolates that had the same sequences throughout the target recombination region, we confirmed the identity of the two sequences using assembled contigs of the two isolates.

### Positive selection

We investigated positive selection across the pan genome of the two major resistant clones to identify genes that could be associated with the spread; the GPSC904;9 isolates from IPD cases (n = 114) and the GPSC932 isolates from IPD cases (n = 92). We did not include non-IPD isolates in this analysis to prevent sampling bias. In each analysis, we determined homologue genes using Roary. The gene clusters that had four or more gene copies were used in this analysis, as RAxML requires four or more operational taxonomic unit sequences to construct a phylogenetic tree. Using these trees and aligned nucleotide sequences of the clusters, we performed a branch-site unrestricted statistical test for episodic diversification (BUSTED) as implemented in HyPhy^30, 31^. In this analysis, we assessed for recombination using the pairwise homoplasy index (PHI) as implemented in Phi^32^ and analyzed only recombination-free genes at each statistical step while adjusting the *P* value following the FDR procedure of Benjamini and Hochberg^33^ to account for increased type I errors due to multiple hypothesis testing. The statistical tests were considered significant when the adjusted *P* value fell below the FDR threshold of 0.05. The selected genes were re-annotated using blastx (https://blast.ncbi.nlm.nih.gov/Blast.cgi; final access date: 9/1/2022). Gene Ontology (GO) terms were assigned to the top hit proteins using Interproscan v5.51-85.0^34^

### Identifying the origin of *pbp1a*-13 using isolates collected in 1996

We subjected 57 PCG isolates with MIC ≥ 0.25 that were collected in Japan in 1996 and stocked in Antimicrobial Resistance Research Center, National Institute of Infectious Diseases to whole-genome sequencing analysis to identify the origin of *pbp1a*-13. These isolates were collected from five regions that covered the entire Japan (Hokkaido, Tohoku, Kanto, Kansai, and Kyusyu region) from adult (n = 35) and pediatric patients (n = 22) with IPD or non-IPD.

## Results

Through whole genome sequencing, we obtained good quality raw reads for 1305 of 1358 tested isolates collected between 2012 and 2017; thus, we conducted subsequent analysis for the 1305 isolates. The average (±SD) number of contigs was 49.0 (±9.8) and N50 (shortest contig length needed to cover 50% of the genome) was 75,565 (±18,877) (Table S1).

### Drug resistant clusters and PBP profile in a core genome phylogenetic tree

We constructed a core genome ML tree using core genome SNPs from the 1305 isolates collected in Japan between 2012 and 2017. We discovered 44 GPSCs and 103 isolates that were not assigned any GPSCs. Among the 44 GPSCs, the most prevalent one was GPSC904;9 (n = 206) followed by GPSC106 (n = 188) and GPSC932 (n = 144). Serotypes and sequence types (STs) associated with each GPSC were summarized in Figure 1 and Table S1. In brief, the most prevalent GPSC904;9 included 196 (95.1%) 15A-CC63 isolates, GPSC106 included 186 (98.9%) 24B/F-CC2572 isolates, and GPSC932 included 144 (100%) 19A-CC3111 isolates. Among the not-assigned isolates, the most prevalent clone was 10A-ST5236 (n = 29) followed by 38-ST6429 (n = 19) and 6C-ST2924 (n = 8). Of the 1305 isolates, 560 (42.9%), 208 (15.9%), and 253 (19.4%) isolates were resistant to PCG (i.e., PG-R), CTX (i.e., CTX-IR and -R), and MEM (i.e., MEM-IR and -R), respectively (Figure 2). In the PG-R, CTX-IR or -R, and MEM-IR or -R isolates, the largest cluster was GPSC904;9 (15A-CC63). We discovered 317 Mβ-lacR isolates (24.3%), the largest Mβ-lacR GPSC being GPSC904;9 (15A-CC63, n = 144) followed by GPSC59 (35B-CC558, n = 58), GPSC932 (19A-ST3111, n = 32), and GPSC4 (mainly 15B/C-CC199; n = 18).

**Figure 1.**
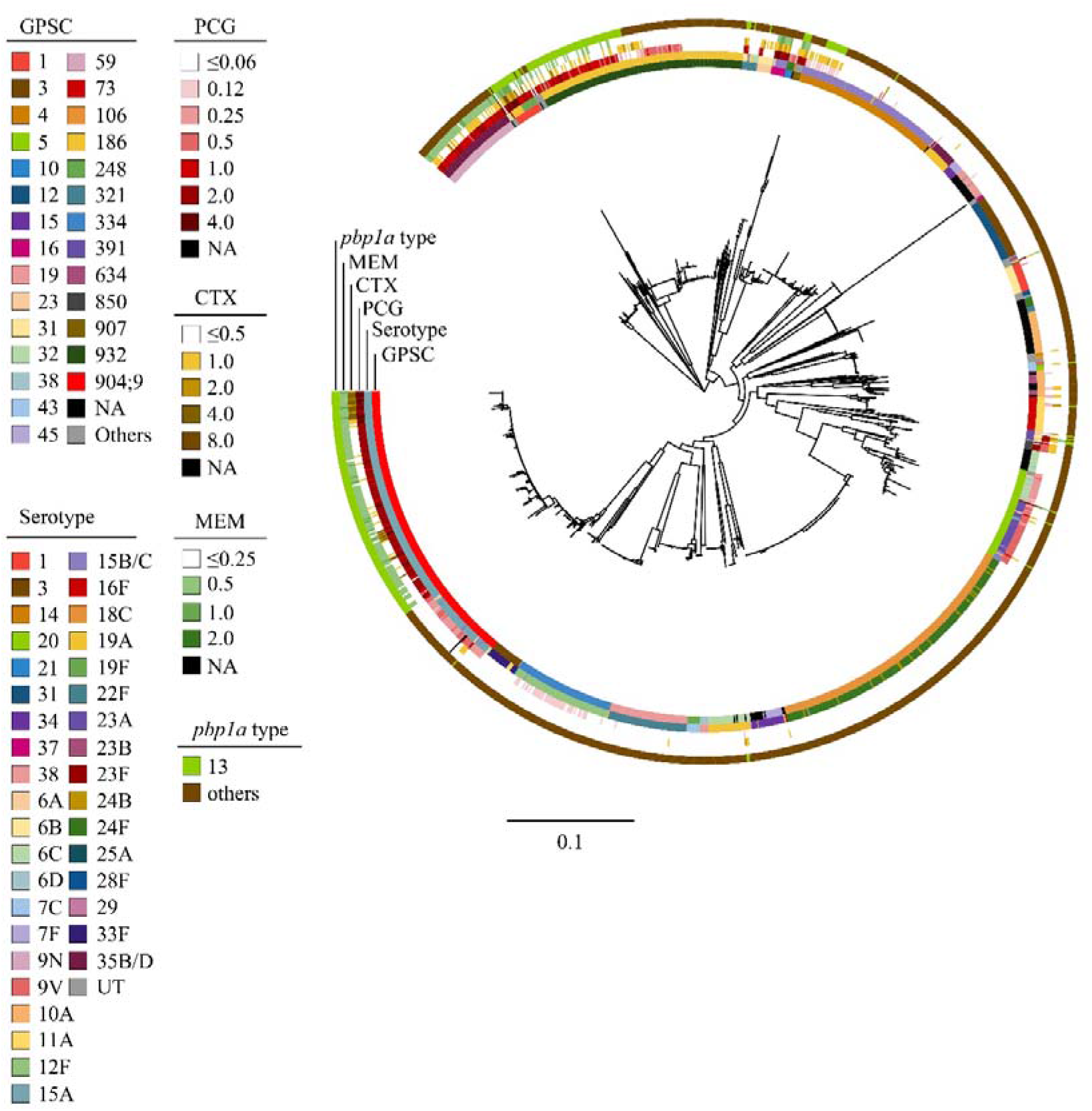
Core-genome based maximum likelihood tree of 1305 pneumococcal isolates collected in Japan, 2012-2014. MICs of each antibiotic, serotype, and GPSC are shown in outside the tree. MIC was measured in mg/l. NA, not assigned; UT, untypeable; PCG, penicillin G; CTX, cefotaxime; MEM, meropenem; GPSC, Global Pneumococcal Sequence Cluster; MIC, minimum inhibitory concentration.

**Figure 2.**
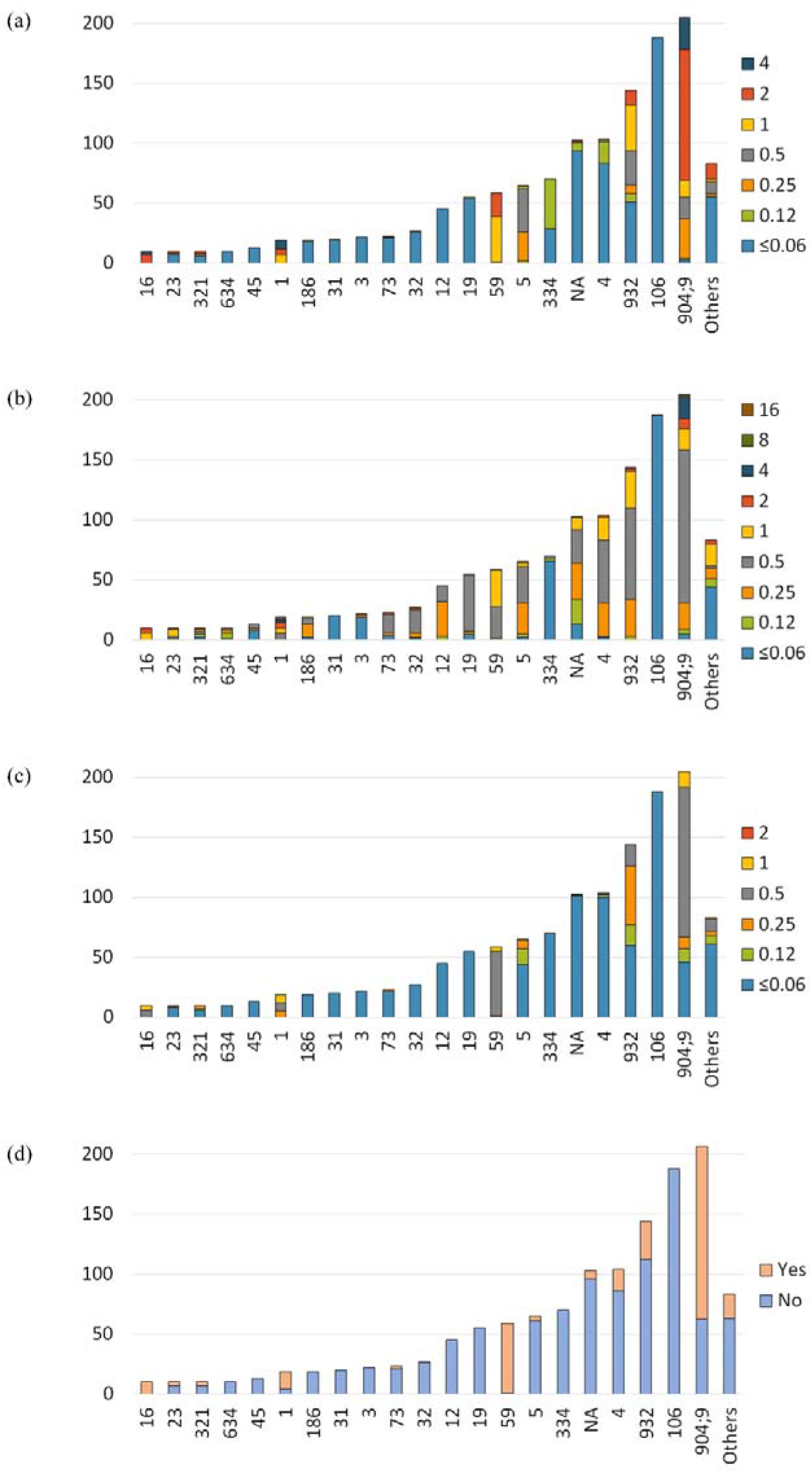
The MIC distributions of PCG, CTX, MEM, and multi β-lactam resistance in each GPSCs. Others include GPSC6, 7, 10, 14, 15, 35, 38, 43, 57, 60, 67, 89, 123, 142, 191, 230, 248, 324, 382, 391, 397, 777, 850, and 904 of which the number of isolates was <10. NA, not assigned. (a) The MIC distribution of PCG (mg/l) in each GPSC. (b) The MIC distribution of CTX (mg/l) in each GPSC. (c) The MIC distribution of MEM (mg/l) in each GPSC. (d) The distribution of multi β-lactam resistance.

Among 560 PG-R isolates, 260 (46.4%) isolates had *pbp1a*-13 and when limiting high-level penicillin resistance isolates with PCG-MIC ≥ 2.0, 175/210 (83.3%) had *pbp1a*-13 (Figure 3). Similarly, 81.4% (48/59) and 73.3% (23/30) of CTX-R and MEM-R isolates had *pbp1a*-13, respectively. In 317 Mβ-lacR isolates, 213 (67.2%) had *pbp1a*-13. We did not observe any relationship between *pbp2b* and *pbp2x* profiles and drug resistance. For instance, the most prevalent *pbp2b* and *pbp2x* types in PG-R isolates were *pbp2b*-175 (n = 145, 25.9%) and *pbp2x-43* (n = 154, 27,5%) as the most prevalent Mβ-lacR GPSC904;9 (15A-CC63) had these types. A total of 273 isolates (20.9%) had *pbp1a*-13, which was the second most prevalent *pbp1a* type, while the most prevalent *pbp1a* type was *pbp1a-2*. Among 273 *pbp1a*-13 positive isolates, 260 (95.2%), 118 (43.2%), and 180 (65.9%) isolates were PG-R, CTX-IR or -R, and MEM-IR or -R, respectively (Figure 4).

**Figure 3.**
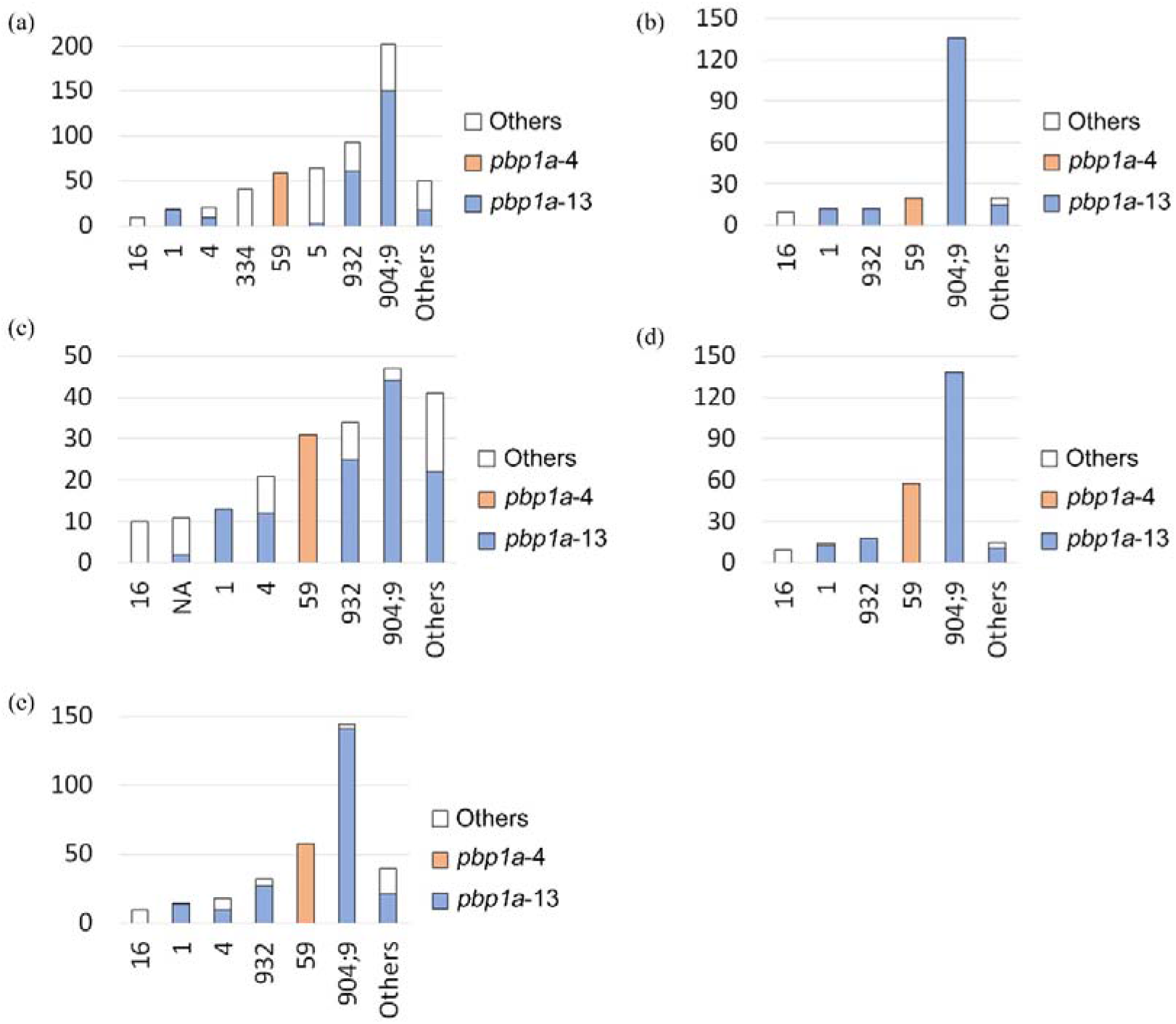
The prevalence of *pbp1a*-13 and −4 in PCG, CTX, MEM, and multi β-lactam resistant GPSCs. NA, not assigned. (a) The prevalence of *pbp1a*-13 and −4 in PCG resistant GPSCs. Other GPSCs include GPSC6, 10, 14, 19, 23, 31, 32, 73, 89, 186, 321, 382, 391, 777, 850, 907, and not assigned. (b) The prevalence of *pbp1a*-13 and −4 in PCG high-level resistant GPSCs. Other GPSCs include GPSC5, 6, 14, 23, 321, 391, 777, 907, and not assigned. (c) The prevalence of *pbp1a*-13 and −4 in CTX resistant GPSCs. Other GPSCs include GPSC5, 6, 14, 19, 23, 32, 43, 73, 186, 321, 391, 777, 850, and 907. (d) The prevalence of *pbp1a*-13 and −4 in MEM resistant GPSCs. Other GPSCs include GPSC4, 5, 6, 14, 391, 777, 907, and not assigned. (e) The prevalence of *pbp1a*-13 and −4 in multi β-lactam resistant GPSCs. Other GPSCs include GPSC5, 6, 14, 23, 32, 73, 321, 777, 850, 907, and not assigned.

**Figure 4.**
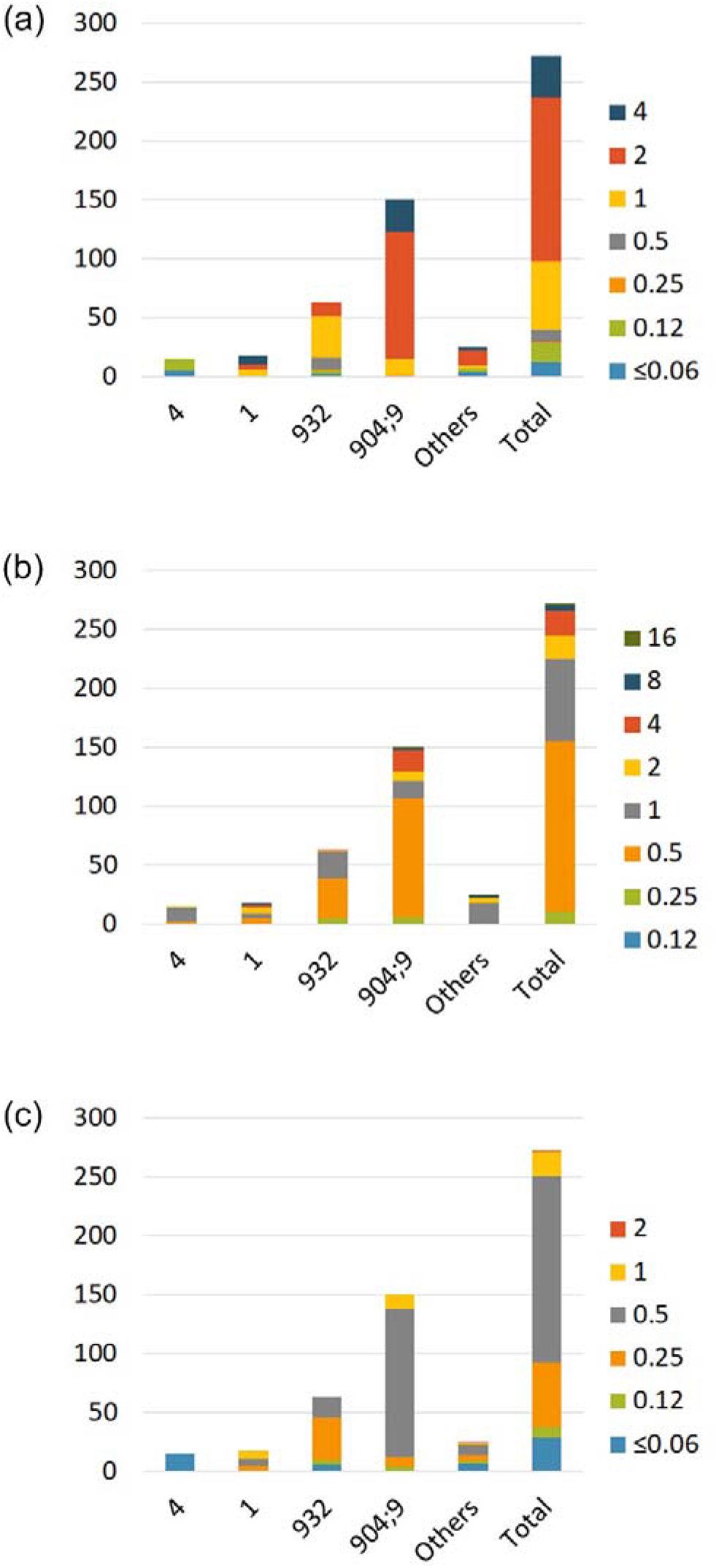
The MIC distributions of PCG, CTX, and MEM in *pbp1a*-13 positive isolates. Other GPSCs include GPSC4, 14, 23, 32, 43, 73, 321, 391, 907, and not assigned. (a) The MIC distribution of PCG (mg/l) in each GPSC. (b) The MIC distribution of CTX (mg/l) in each GPSC. (c) The MIC distribution of MEM (mg/l) in each GPSC.

Furthermore, we observed a high prevalence of the macrolide resistance genes *ermB* and *mefE*. The *ermB* and *mefE* were detected in 1035/1305 (79.3%) and 355/1305 (27.2%) isolates, respectively, while 178/1305 (13.6%) isolates had both genes (Figure 5). The distribution of these macrolide resistance genes was fundamentally related to GPSC and serotype. For instance, 141/144 (97.9%) GPSC 932 (19A-CC3111) isolates had both *ermB* and *mefE* genes, while all 206 GPSC904;9 (15A-CC63) isolates had only *ermB*. However, we discovered GPSCs that showed diverse prevalence of these genes in GPSC19 (22F-CC433), a cluster that contained 33 isolates with only *ermB*, eight isolates with only *mefE*, and 14 isolates without any macrolide resistance genes while the diversity of the cluster was not high (π and Θ showed similar values as other clusters [see the results below]). This may indicate that this cluster tended to transfer *mefE-* and/or *ermB*-possessing transposons (i.e., Tn*916*-like integrative conjugative element) via an unknown mechanism. A total of 125 isolates (9.6%) and 428 isolates (32.8%) had *folA* I100L mutation and *folP* insertion, respectively. The largest GPSCs with the *folA* I100L mutation and *folP* insertion were GPSC334 (12F-CC4846) and GPSC106 (24B/F-CC2572), respectively. Further, we identified 311 MDR isolates (23.8%) and similar to Mβ-lacR, GPSC904;9 was the largest cluster (n = 144), followed by GPSC59 (35B-CC558, n = 54), GPSC932 (19A-CC3111, n = 32), and GPSC4 (15B/C-CC199, n = 18).

**Figure 5.**
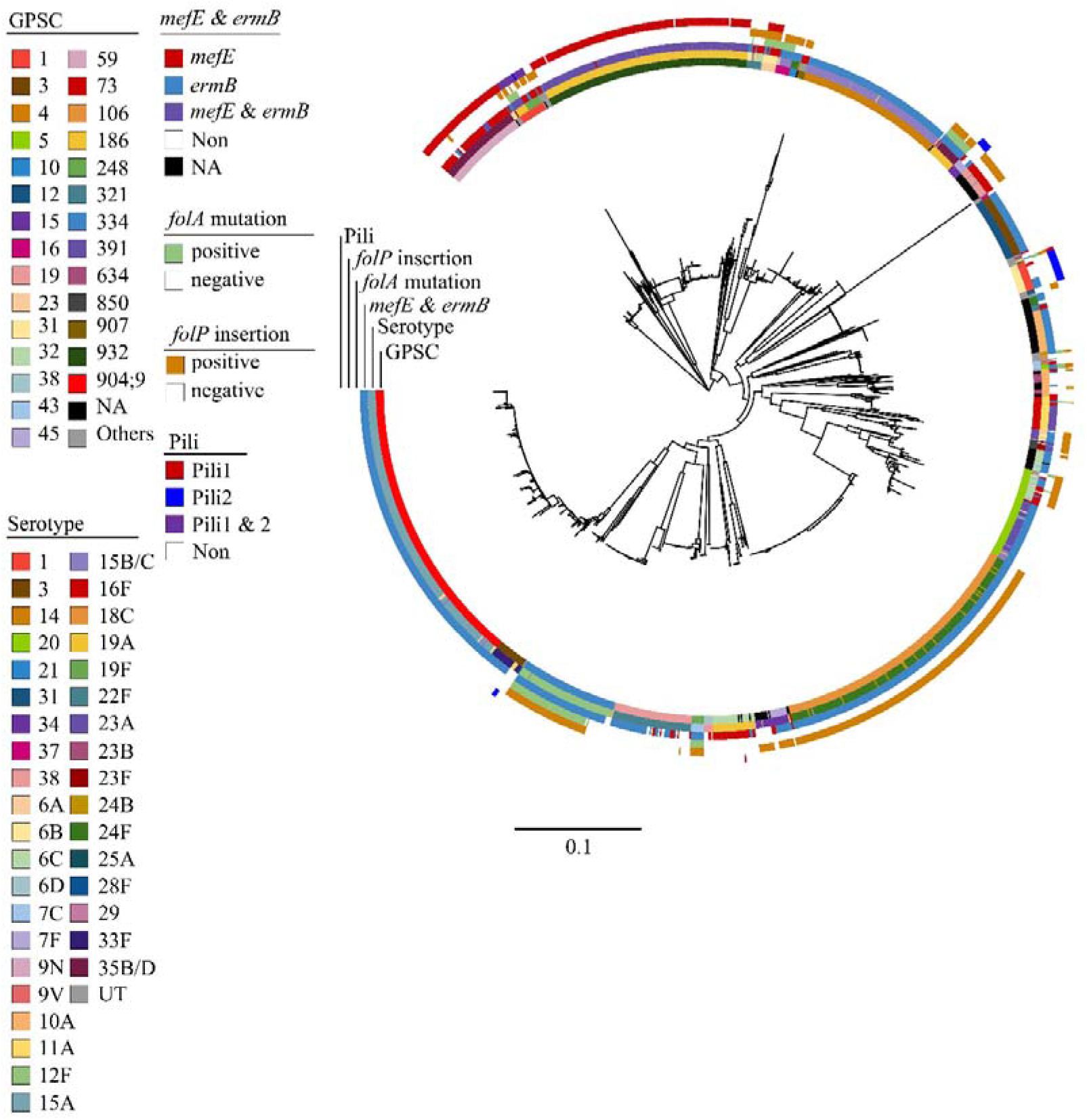
Core-genome based maximum likelihood tree of 1305 pneumococcal isolates collected in Japan, 2012-2014. The presence of resistance genes (*ermB, mefE, folA* mutation, and *folP* insertion), pili (Pili1 and Pili2), serotype, and GPSC are shown in the outside the tree. NA, not assigned; UT, untypeable.

### Genetic diversity and population dynamics of pneumococci collected from Japan

We investigated the genetic diversity and population dynamics of each GPSC with 10 or more IPD isolates (Table 1) to prevent sampling bias. GPSC4 (serotype 15B/C-CC199), GPSC19 (serotype 22F-CC433), GPSC106 (24B/F-CC2572), GPSC186 (serotype 35B-CC2755), GPSC634 (serotype 10A-CC1263), GPSC904;9 (serotype 15A-CC63), and GPSC932 (serotype 19A-CC3111) exhibited significant negative value for Tajima’s D, Fu and Li’s D*, and Fu and Li’s F*, demonstrating a deviation from neutral effects. All populations exhibiting this deviation consisted of non-PCV13 serotype isolates except for GPSC932, which consisted of serotype 19A-CC3111 isolates that increased in a number of IPD cases after the introduction of PCV7 in Japan. Conversely, GPSC1 (serotype 19A/F-CC236/320), GPSC23 (serotype 6B-CC2983), and GPSC31 (serotype 1-CC306) that were covered by PCV13 did not show deviations from neutrality.

The theta of each population ranged between 0.00004 and 0.00084. The GPSC3 (serotype 33F-CC717) theta was the highest (0.00084) and more than three times higher than that of GPSC634 (serotype 10A-CC1263; 0.00027), which was the second highest. This indicates that GPSC3 had considerably more molecular diversity than any other GPSC.

### Divergence dating analysis suggested *pbp1a*-13 was inserted into major GPSCs before the introduction of PCVs

We estimated the date when *pbp1a*-13 was introduced into the two major resistant GPSCs, GPSC904;9 (serotype 15A-CC63) and GPSC932 (serotype 19A-CC3111) (Figure 6 and 7). Maximum clade credibility trees of the two GPSCs showed that *pbp1a*-13 appeared to be recombined into GPSC904;9 and GPSC932 via recombination between 1994.4 (95% highest posterior density [HPD]: 1989.6-1998.4) and 1997.2 (95% HPD:1993.0-2000.6) as well as between 2001.2 (95% HPD: 1999.7-2002.9) and 2005.8 (95% HPD: 2005.1-2006.4).

**Figure 6.**
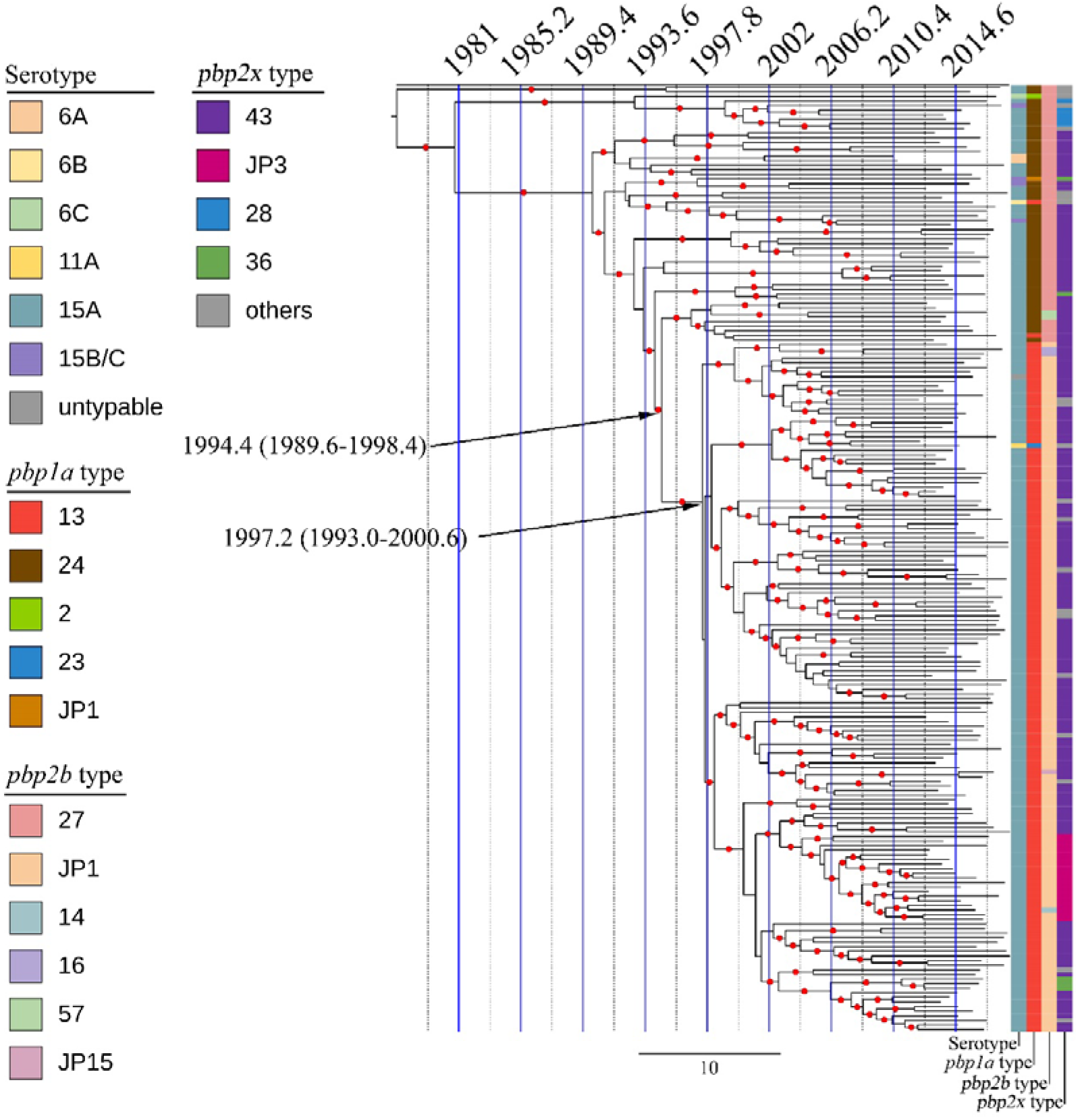
Maximum-clade credibility tree of GPSC904;9 isolates. Red dots indicate that the branches were supported with posterior values ≥0.7. The estimated times between the introduction of *pbp1a*-13 into GPSC904;9 are shown on the tree with 95% highest posterior density.

**Figure 7.**
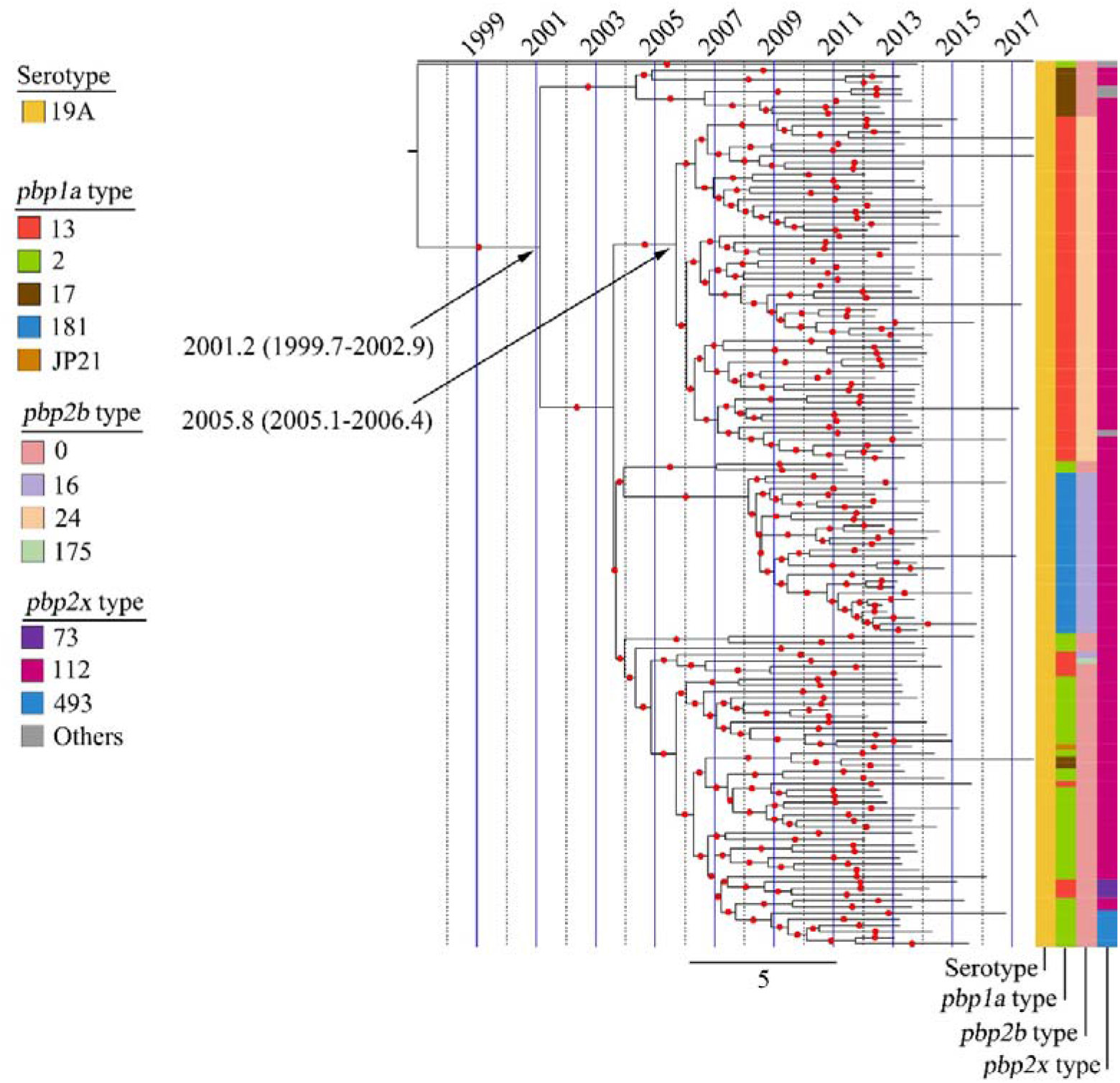
Maximum-clade credibility tree of GPSC932 isolates. Red dots indicate that the branches were supported with posterior values ≥0.7. The estimated times between the introduction of *pbp1a*-13 into GPSC932 are shown on the tree with 95% highest posterior density.

### Evidence that *pbp1a*-13 was inserted into different GPSCs following recombination events

Our divergence time analysis suggests that *pbp1a*-13 was inserted into two resistance clusters, GPSC904;9 and GPSC932, at difference times before the introduction of PCVs. These results indicated that the two resistance clusters arouse independent of the introduction of PCVs and the pressure of PCVs may increase the population to be detected in clinical settings as the serotypes included in the clusters (i.e., serotype 15A and 19A) were not included in the PCV7. Alternatively, we identified isolates with *pbp1a*-13 that did not form clusters in GPSC4, 5, 23, 32, 43, 321, 73, 391, and 904;9; as the isolates did not spread with clonality. In such cases, the recombination events that inserted *pbp1a*-13 into isolates might have occurred recently (between the final node and the leave) and the sequence might be conserved; thus, we investigated the estimated recombination regions that included *pbp1a*-13 to validate the possibility that recombination events cause the spread of *pbp1a*-13 between different GPSCs. By comparing recombination regions that included *pbp1a*-13 regions to regions of other isolates, we uncovered two cases that had a *pbp1a*-13-containing recombination region whose nucleotide sequence was identical to that of another isolate included in a different GPSC. PC0036 (GPSC321, serotype 6B-ST2923) with *pbp1a*-13 had a 15,234-bp recombination region (Supplementary figure 1 and 2) and the nucleotide sequence of the region was identical to that of a corresponding region of PC0362 (GPSC5, serotype 6A-ST338) and PC0329 (GPSC14, serotype 23F-ST242). Similarly, PC0070 (GPSC391, serotype 6B-ST2756) with *pbp1a*-13 had a 4188-bp recombination region (Supplementary figure 3 and 4) and the nucleotide sequence of the region was identical to that of a corresponding region of PC0036 (GPSC321, serotype 6B-ST2923), PC0299 (GPSC23, serotype 6B-ST5497), PC0329 (GPSC14, serotype 23F-ST242), PC0362 (GPSC5, serotype 6A, ST338), and PC0442 (GPSC32, serotype 19A-ST5237).

### Estimation of the origin of *pbp1a*-13 using isolates from 1996

To estimate the origin of *pbp1a*-13, which was imported into major resistant clones in Japan (GPSC904;9 and GPSC932), we subjected 57 pneumococcal isolates collected in 1996 in Japan to whole-genome sequencing. These isolates comprised of 17 serotypes (serotype 3, 6B, 6C, 9A, 9V, 10A, 11A, 12F, 13, 14, 15A, 19F, 23A, 23F, 24F, 35B, and 37) and the most prevalent GPSC was GPSC1 (serotype 19F-CC236, n = 17) followed by GPSC12 (serotype 3-ST180, n = 12) and GPSC14 (serotype 23F-CC242, n = 9) (Table S2). Among these isolates, we discovered 20 isolates that possessed *pbp1a-13:* 14 GPSC1, serotype 19F-ST236, five GPSC14 serotype 23F-CC242, and one GPSC-not-assigned, serotype 23F-new ST isolate. There were no GPSC904;9 and GPSC932 isolates in this collection.

### Positive selection

We assessed positive selection across the pan genome of the GPSC904;9 and GPSC932 datasets using only IPD isolates. We identified seven genes (three core genes and four accessory genes) in GPSC904;9 pan genome and seven genes (one core gene and six accessory genes) in GPSC932 pan genome that were significant for positive selection (Table S3 and S4). In the GPSC904;9 dataset, PBP1A was positively selected (adjusted *P* = 1.64E^-5^), while in the GPSC932 dataset, PBP1A was determined as recombinant and was not subjected to the selection test. In the GPSC932 dataset, StrH, surface-anchored exoglycosidases that was one of the LPXTG-anchored surface proteins, was positively selected (adjusted *P* = 0.027)

## Discussion

We analyzed 1305 pneumococcal isolates collected over a six year period throughout Japan after the introduction of PCVs. We demonstrated that *pbp1a*-13 was highly prevalent in β-lactam resistant pneumococci in Japan; 83.3%, 81.4%, and 73.3% of high-level penicillin resistant, CTX-R, and MEM-R isolates had *pbp1a*-13. In Japan, the two major β-lactam-resistant clones were GPSC904;9 and GPSC932, which were comprised mainly of serotype 15A-CC63 and serotype19A-CC3111, respectively, while Mβ-lacR isolates distributed in a total of 18 GPSCs. We discovered *pbp1a*-13 in 12 of the 18 GPSCs in Japan; thus, suggesting the possibility that this *pbp1a*-13 transferred horizontally between different GPSCs resulting in the emergence and the spread of resistant GPSCs. PBP1a-13 had a 370SSMK substitution in their β-lactam binding SXXK motif, which has been shown to cause β-lactam-resistant including PCG with PBP1B and/or PBP2X alterations^35, 36^. In our collection, MICs of PCG among isolates with *pbp1a*-13 ranged between 0.06-4 μg/ml (CTX, 0.12-16; MEM, ≤0.06-2), therefore, the acquisition of *pbp1a*-13 may not cause β-lactam resistance by itself. However, as Chen et al. demonstrated in a previous study, the introduction of *pbp1a*-13 increased MICs of β-lactams with alterations of PBP2B and/or PBP2x^35^. Therefore, we hypothesize that the spread of *pbp1a*-13 should be considered a risk factor for the emergence of novel β-lactam resistance clones. Hence, we revealed two cases where the exact same sequence of the recombination region including pbp1a and its surroundings was present in the isolates of another GPSC, indicating that the horizontal transmission of *pbp1a*-13 occurred.

We estimated the date when *pbp1a*-13 was transmitted into two major β-lactam resistant GPSCs in Japan (GPSC904;9 and GPSC932) using the Bayesian Markov Chain Mote Carlo framework. Our analysis indicates that *pbp1a*-13 was transmitted into the GPSC904;9 between 1994 and 1997 and into GPSC932 between 2001 and 2006. Furthermore, we confirmed the presence of *pbp1a*-13 at an earlier time (1994-1997) using the 1996 pneumococcal collection isolates stocked from the National Institute of Infectious Diseases in Japan. We obtained draft genome data of all of available 57 isolates collected in 1996. Consequently, we identified 20 isolates with *pbp1a*-13 including 14 GPSC1 (serotype 19F-ST236) and five GPSC14 (serotype 23F-CC242) isolates, but no GPSC904;9 and GPSC932 isolates. GPSC1 and GPSC14 were known as PMEN14 (Taiwan^19F^-14) and PMEN15 (Taiwan^23F^-15), respectively. According to the GPSC database (final access: 2022. October 6^th^), 210 of the 229 GPSC1 isolated with serotype 19F-ST236 possess *pbp1a*-13 and were distributed across 16 countries as detected between 1997 and 2015. Similarly, 31 of the 85 GPSC14 isolates with serotype 23F-ST242 possess *pbp1a*-13 and were distributed across 7 countries as detected between 2000 and 2016. Therefore, these two GPSCs containing *pbp1a*-13 appear not to be minor strains before and after the introduction of PCV7. Considering these results, we concluded that (1) *pbp1a*-13 was transmitted to GPSC904;9 and GPSC932 before the introduction of PCVs in Japan (PCV7 was licensed in use in 2012 in Japan) from isolates with *pbp1a*-13 such as serotype 19F-ST236 and serotype 23F-CC242 that were prevalent at that time; however, GPSC904;9 and GPSC932 were not detectable due to the low prevalence. (2) After the introduction of PCVs, GPSC904;9 and GPSC932 were selected and increased to detectable levels due to the vaccines pressures, as they were not covered by PCV7 and/or PCV13.

We have to note the risk of emergence and increase in β-lactam resistant lineages within Japan because isolates with *pbp1a*-13 could change to Mβ-lacR through the acquisition of altered *pbp2b* and/or *pbp2x*, while *pbp1a*-13 also appears to be transmittable. Moreover, in the GPSC904;9 population, *pbp1a* was under positive selection. While we have to be careful with the selection test results interpretation as the existence of recombination increases the false positive in this analysis^37^ and *S. pneumoniae* is a transformable pathogen by natural competence^38^, we have to note the possibility that this clone is positively selected for *pbp1a*-13 due to the use of antibiotics.

Neutrality tests showed that GPSCs whose dominant serotypes were not covered by PCVs significantly were deviated from neutrality. While negative values of Tajima’s D, Fu & Li’s F*, and D* are also caused by directional selection, our result could indicate that the expansion of PCVs-uncovering population occurred under the pressure of PCVs. Of note, although GPSC1 (serotype 19A/F-CC236/320) and GPSC932 (serotype 19A-CC3111) had the same dominant serotype (19A/F) that is covered by PCV13, only GPSC932 deviated from neutrality. After the introduction of PCV7, the most dominant clone in serotype 19A in Japan was CC3111 while it was CC236/320 in several other countries^5,24, 39, 40, 41^; therefore, GPSC932 (serotype 19A-CC3111) isolates may exhibit fitness factors that allow them to thrive in Japan’s circumstances and could outperform GPSC1 isolates. Although the fitness factors remain unclear, we have to note that the resistance rate to MEM of GPSC932 was lower than that of GPSC1 isolates (12.5% vs 73.4%); therefore, drug resistance was not necessary to fitness. In addition, in our data, only GPSC3 (serotype 33F-CC717) had positive values for neutrality tests as well as high values for nucleotide diversity and theta; thus, suggesting that this GPSC was likely to be an old and not-expanding population. However, according to previous studies in Japan in the 2000’ investigating serotype prevalence, the GPSC3 like-serotype 33F was not observed^42, 43^. Only one study by Nagai et al. demonstrated that one serogroup of 33 isolate was found among 116 isolates that were collected in a regional surveillance study in Japan between 1993 and 1995^44^ In contrast, according to a previous study^45^, GPSC3 with serotype 33F isolates were already prevalent in 2007-2011 in Canada. Therefore, we hypothesize that this cluster was imported into Japan from foreign countries before 1990s and kept stable in the Japanese population, or imported into Japan sporadically at several times after becoming stable in the original regions.

We have to note the limitations of this study. First, we could not analyze some GPSCs due to the relatively small sample size. Second, we could not show the direct evidence that *pbp1a*-13 was introduced into two major resistant GPSCs in Japan (GPSC904;9 and GPSC932) because the introduction occurred in the past. Third, this study was conducted using isolates from Japan where population inflows and outflows are relatively reduced; therefore, it is unclear if the concept of *pbp1a* transmission could be placed into a global context. However, this is the first study that showed the process of drug-resistant pneumococcal lineages epidemiology at a nationwide scale. Furthermore, we consider *pbp1a* (i.e., *pbp1a*-13) transmission as a potential risk factor for the emergence and spread of resistance lineages. While amino acid alterations in one PBP of PBP1A, PBP2B, and PBP2X may cause low-level resistance to β-lactams, the broad distribution of that kind of PBP via recombinational events may lead the emergence of high-level and/or multi-drug resistant lineages due to the acquisition of additional PBP alterations. This study also underlines the need for the pneumococcal molecular epidemiology studies to monitor the trend of PBP type underlying serotype changes for the early detection of emerging β-lactam resistance lineages. In addition, the recognition of such risks can guide proper and timely policy making regarding the choice of PCV to prevent the emergence and the spread of β-lactam resistant pneumococci.

## Author contribution

T.F and S.S designed and coordinated the surveillance study. S.N and B.C performed the experiments. S.N. analyzed the experimental data. S.N wrote the manuscript. Y.I, Y.M, M.Y, M.N, M.S, and M.S supervised the experiments.

## Acknowledgement

This study was supported by AMED grant number JP21fk0108147 and by JSPS KAKENHI under grants number 19K16637 and 21K15432 (S.N). This work was supported by the Japan AMED grant number JP21fk0108139 and JP21fk0108099 (B.C).

