## Supplementary material for "Horizontal transmission of penicillin binding protein 1A caused a nationwide spread of β-lactam resistance in pneumococci"

Supplementary Figures

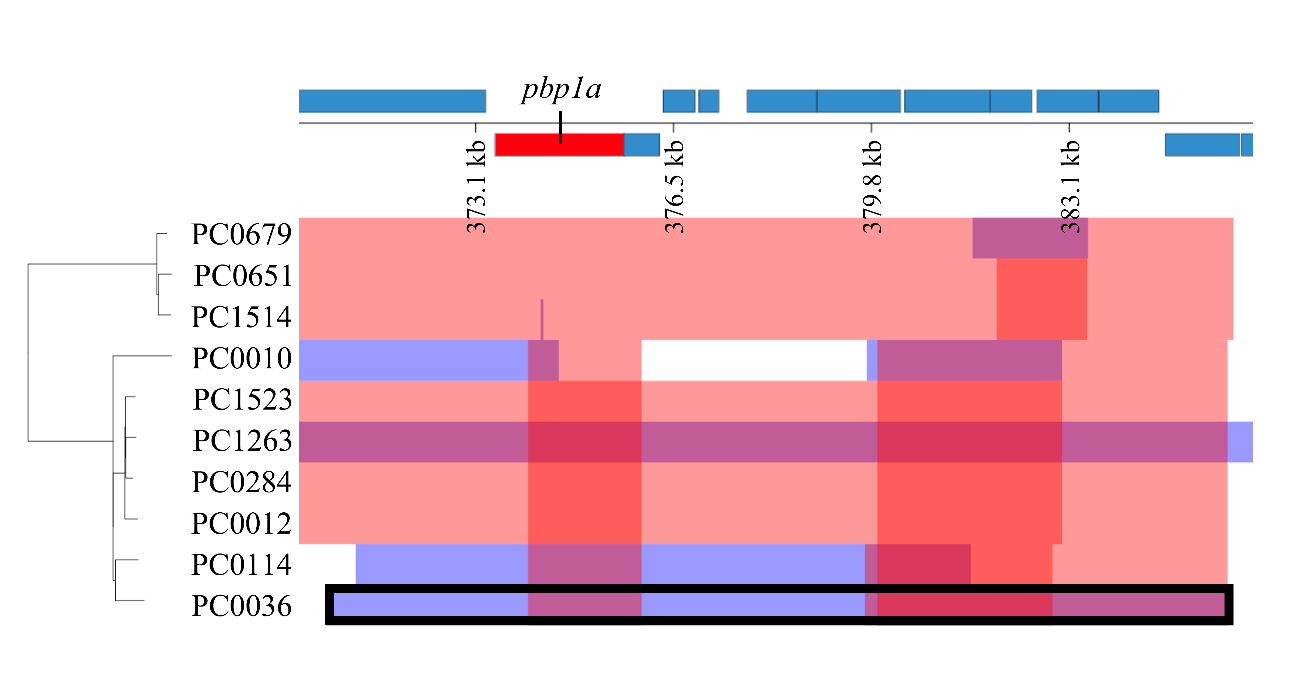

Figure S1. The recombination site detected in PC0036 (GPSC321, serotype 6B-ST2923) using Gubbins is shown with a black contour. This region had a 15,324-bp length overlapping with *pbp1a*. The red bars indicate recombinations reconstructed as occurring on internal branches, which are therefore shared by isolates through common ancestry. The blue bars indicate recombinations reconstructed as occurring on terminal branches, which are unique to individual isolates.

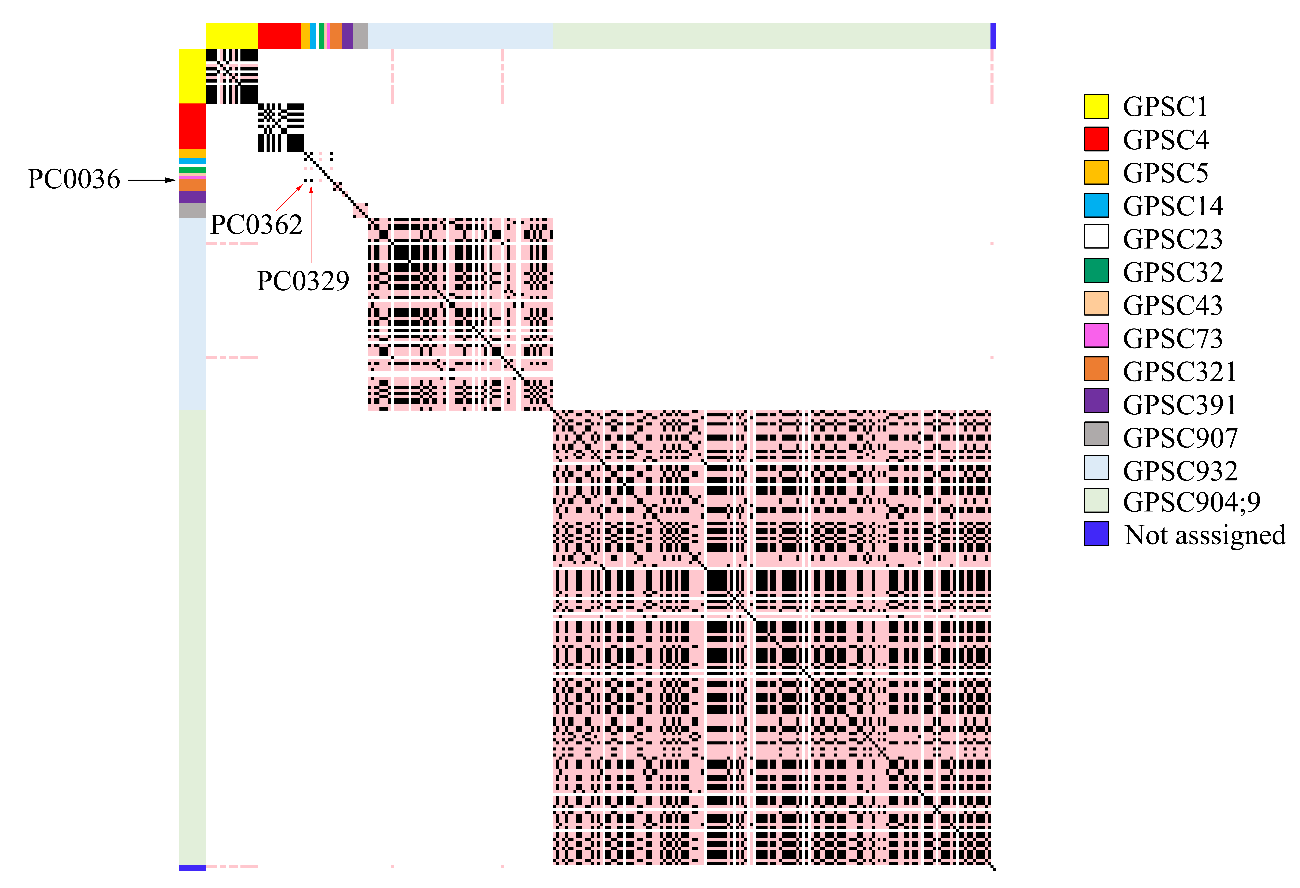

Figure S2. Heatmap of the number of SNPs detected from NGS reads of each isolate with *pbp1a*-13 (n = 273) and mapped to the 15,324-bp recombination region of PC0036 that was estimated to have imported *pbp1a*-13 into PC0036. Black cells indicate the corresponding two isolates had no SNP and pink cells indicate they had 1-5 SNPs. PC0036 (GPSC321, serotype 6B-ST2923) contained an identical nucleotide sequence of the region to those of PC0362 (GPSC5, serotype 6A-ST338) and PC0329 (GPSC14, serotype 23F-ST242).

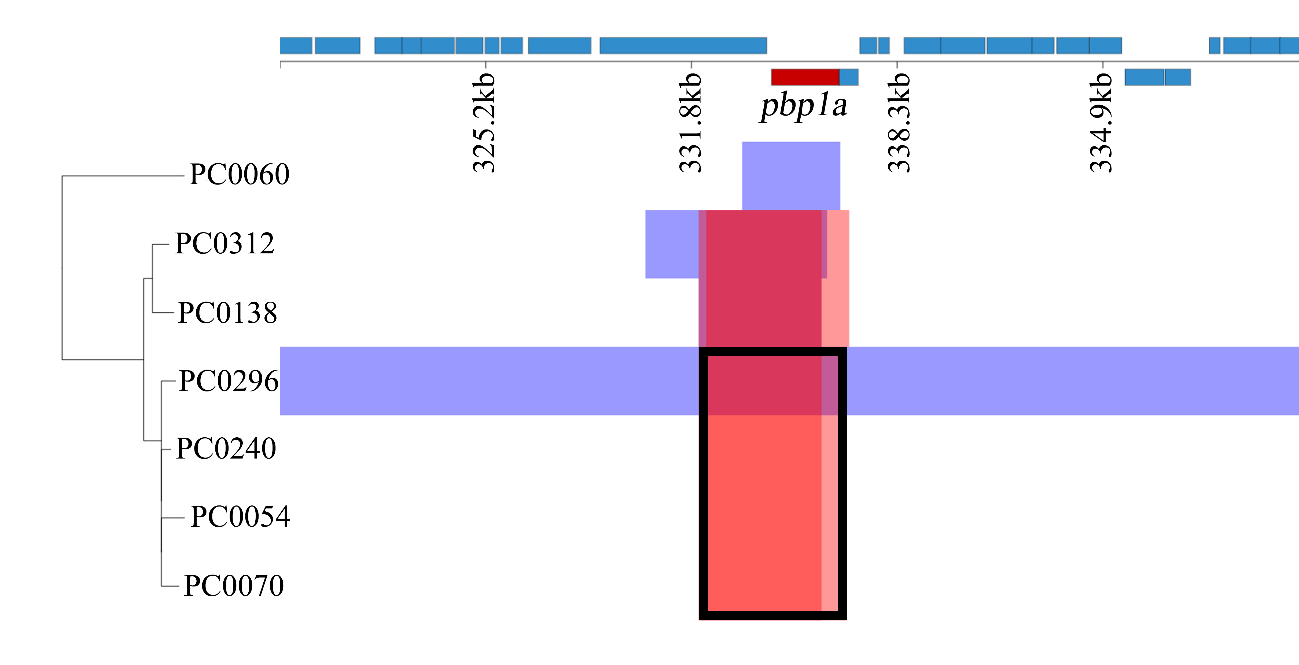

Figure S3. The recombination site detected in PC0070 (GPSC391, serotype 6B-ST2756) using Gubbins is shown with a black contour. This region had a 4,188-bp length overlapping with *pbp1a*. The red bars indicate recombinations reconstructed as occurring on internal branches, which are therefore shared by isolates through common ancestry. The blue bars indicate recombinations reconstructed as occurring on terminal branches, which are unique to individual isolates.

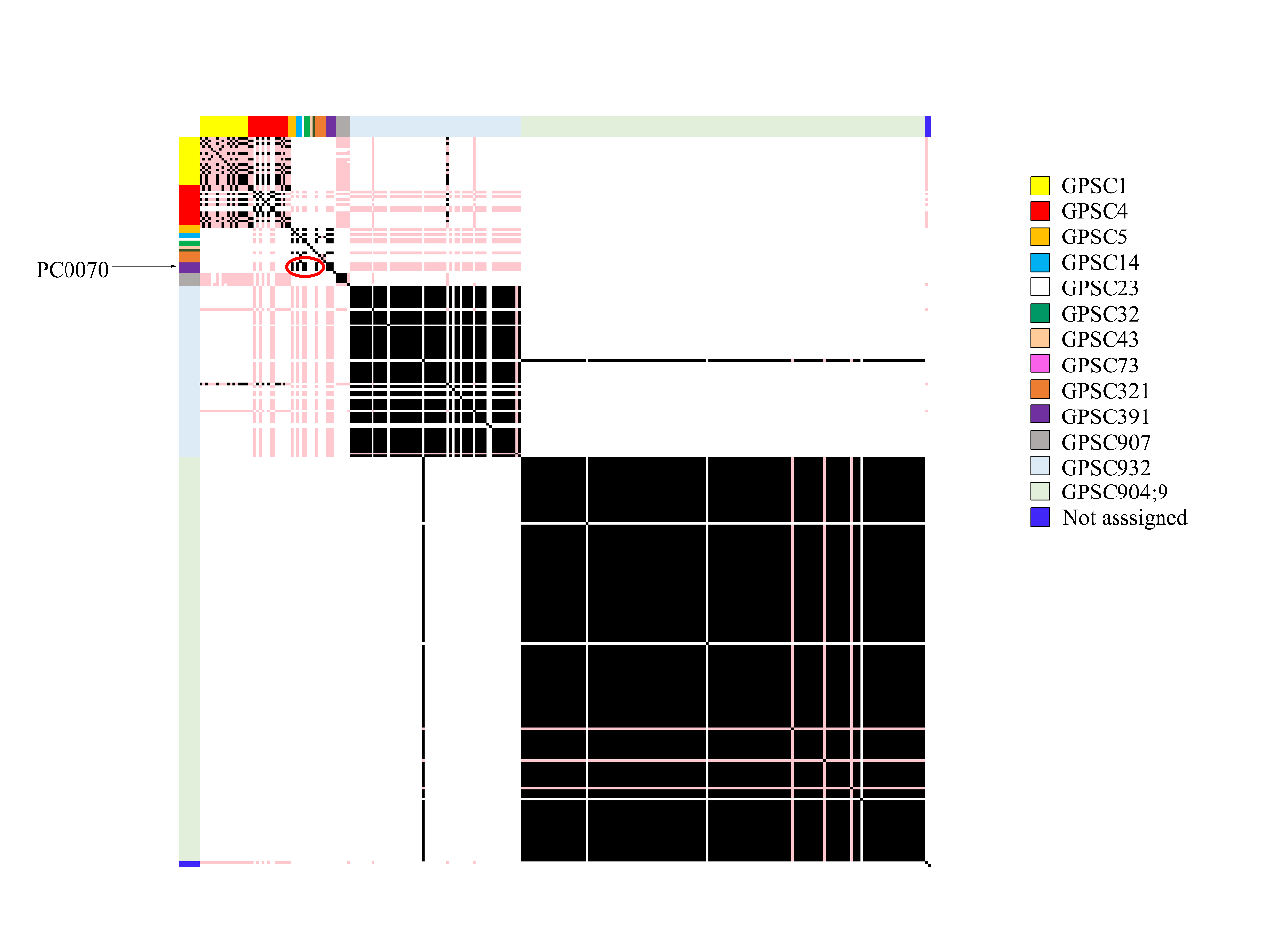

Figure S4. Heatmap of the number of SNPs detected from NGS reads of each isolate with *pbp1a*-13 (n = 273) and mapped to the 4,188-bp recombination region of PC0070 that was estimated to have imported *pbp1a*-13 into PC0070. Black cells indicate the corresponding two isolates had no SNPs and pink cells indicate they had 1-5 SNPs. PC0070 (GPSC391, serotype 6B-ST2756) contained an identical nucleotide sequence of the region as those of PC0299 (GPSC23, serotype 6B-ST5497), PC0329 (GPSC14, serotype 23F-ST242), PC0362 (GPSC5, serotype 6A, ST338), and PC0442 (GPSC32, serotype 19A-ST5237), which are circled by a red oval.
